# HPRep: Quantifying reproducibility in HiChIP and PLAC-seq datasets

**DOI:** 10.1101/2020.11.23.394239

**Authors:** Jonathan D. Rosen, Yuchen Yang, Armen Abnousi, Jiawen Chen, Michael Song, Ian R. Jones, Yin Shen, Ming Hu, Yun Li

## Abstract

HiChIP and PLAC-seq are emerging technologies for studying genome-wide long-range chromatin interactions mediated by protein of interest, enabling more sensitive and cost-efficient interrogation of protein-centric chromatin conformation. However, due to the unbalanced read distribution introduced by protein immunoprecipitation, existing reproducibility measures developed for Hi-C data are not appropriate for the analysis of HiChIP and PLAC-seq data.

Here, we present HPRep, a stratified and weighted correlation metric derived from normalized contact counts, to quantify reproducibility in HiChIP and PLAC-seq data. We applied HPRep to multiple real datasets and demonstrate that HPRep outperforms existing reproducibility measures developed for Hi-C data. Specifically, we applied HPRep to H3K4me3 PLAC-seq data from mouse embryonic stem cells and mouse brain tissues, as well as H3K27ac HiChIP data from human lymphoblastoid cell line GM12878 and leukemia cell line K562, showing that HPRep can more clearly separate among pseudo-replicates, real replicates, and non-replicates. Furthermore, in an H3K4me3 PLAC-seq dataset consisting of 11 samples from four human brain cell types, HPRep demonstrates expected clustering of data which could not be achieved by existing methods developed for Hi-C data, highlighting the need of a reproducibility metric tailored to HiChIP and PLAC-seq data.

## 1 Introduction

Chromatin spatial organization plays a critical role in genome structure and transcriptional regulation (Li *et al.*, 2018; Schmitt *et al.*, 2016; Schoenfelder and Fraser, 2019). During the last decade, great strides have been made in the mapping of long-range chromatin interactions, thanks to the rapid development of chromatin conformation capture (3C) based technologies. Among them, Hi-C enables genome-wide measure of chromatin spatial organization and has been widely used in practice. To ensure scientific rigor, various methods have been developed to assess the reproducibility of Hi-C data (Yang *et al.*, 2017; Ursu *et al.*, 2017; Yan *et al.*, 2017; Sauria *et al.*, 2017; Yardimici *et al.*, 2019). For example, HiCRep (Yang *et al.*, 2017) first performs 2D smoothing to reduce the stochastic noise resulting from the sparsity of Hi-C data, and then quantifies reproducibility by calculating a stratified correlation, which is a weighted average of correlation coefficients between contact frequencies across specific one-dimensional (1D) genomic distance bands. HiC-Spector (Yan *et al.*, 2017) adopts a different approach, transforming symmetric Hi-C contact matrices to their corresponding Laplacian matrices and then calculating similarity as the average of the distances between normalized eigenvectors. Similar to HiCRep, GenomeDISCO (Ursu *et al.*, 2017) relies on data smoothing, which is performed over a range of steps of the random walk to determine an optimal separation between biological replicates and non-replicates as measured by area under the precision-recall curve. The reproducibility measure is a function of distances between two contact matrices smoothed using this optimized number of steps. QuASAR-Rep (Sauria *et al.*, 2017) determines a local correlation matrix by comparing observed interaction counts to background signal-distance values within a 100-bin range. This local correlation matrix is transformed by element-wise multiplication with a matrix of scaled interaction counts and the reproducibility between two samples is defined as the Pearson correlation coefficient between the corresponding transformed matrices.

Recently, HiChIP (Mumbach *et al.*, 2016) and PLAC-seq (Fang *et al.*, 2016) technologies (hereafter collectively referred to as HP for brevity) have been developed to study protein-mediated long-range chromatin interactions at much reduced cost and greatly enhanced resolution relative to Hi-C. While the chromatin immunoprecipitation (ChIP) step involved in HP technologies allows for the cost and resolution benefits, it also introduces additional layers of systematic biases which make analysis methods developed for Hi-C data potentially unsuitable for HP data. To date, no method is available for quantifying reproducibility of HP data.

To fill in this gap, we propose a novel method, HPRep, to measure the similarity or reproducibility between two HP datasets. HPRep is motivated by HiCRep (Yang *et al.*, 2017), the previously described method developed for quantifying reproducibility of Hi-C data. Similar to HiCRep, HPRep leverages the dependence of chromatin contact frequency on 1D genomic distance; however, in contrast, HPRep models different ChIP enrichment levels (Supplementary Text S1), which contributes to the systematic biases specific to HP data, and also incorporates an unbalanced data matrix that addresses the targeted structure of HP data in comparison to Hi-C data.

## 2 Methods

Currently available methods to quantify reproducibility in Hi-C datasets, such as HiCRep, HiC-Spector, GenomeDISCO, and QuASAR-Rep (systematically evaluated in Yardimci *et al.*, 2019), all involve derivation of a similarity metric between two contact frequency matrices. The input Hi-C data consists of *n* × *n* symmetric matrices of non-negative integers, where each row/column represents one genomic locus (i.e., bin) and *n* is the total number of bins. The (*i*, *j*) element of such a matrix represents the number of paired-end reads spanning between bin *i* and bin *j*.

These existing methods are conceptually inappropriate for HP data due to the unbalanced read distribution due to ChIP enrichment that is introduced in the HP experiments. In addition, while Hi-C data consist of interactions among all bin pairs, HP data is restricted to bin pairs where at least one bin overlaps a binding region of the protein of interest. Such overlapping bins are referred to as the anchor bins, and two HP datasets may have different sets of anchor bins. We further define bin pairs consisting of two anchor bins as the “AND” pairs, and those consisting of only one anchor bin are defined as the “XOR” pairs. In contrast, the “NOT” pairs, for which neither bin is an anchor bin, are not meaningful due to the nature of HP technologies and therefore are not used in HP data analysis (Juric *et al.*, 2019).

The data structure in HPRep is an *N* × *m* matrix (Supplementary Text S2), where *N* represents the number of anchor bins and *m* = 2 * 1Mb/resolution, where resolution refers to the bin size (1Mb is set as the default but can be modified by the user). The (*i*, *j*) th element represents the normalized contact frequency between anchor *i* and the bin *j* bin widths away from the anchor, *j* ∈ {−*m*/2, …, −1, 1, …, *m*/2}. The number of anchor bins, *N*, is the cardinality of the union set of anchor bins for all datasets in the study. Normalization is performed via a two-step procedure: 1) Raw counts are adjusted for the biases introduced by effective fragment length, GC content, mappability, and ChIP efficiency by fitting a positive Poisson regression model, following the approach detailed in the MAPS method (Juric *et al.*, 2019). Separate models are fit to the AND and XOR pairs since the AND pairs are expected to have significantly higher contact frequencies due to double ChIP enrichment. 2) Using the fitted models, the data are normalized by taking the log2 value of (1 + observed / expected counts). Further details in Supplementary Text S1.

Similar to HiCRep (Yang *et al.*, 2017), the distance metric used by HPRep is a weighted Pearson correlation coefficient that is stratified by distance. Note in **Figure 1** that these strata are the pairs of columns of the previously described data matrix which are equal-distant from the center. Due to the sparsity of HP data, especially for long-range chromatin interactions, the normalized count values are smoothed. The smoothing procedure used is a 1D arithmetic mean of values within a window of *d* bins away along the same row (Supplementary Text S3 for optimization procedure). Each of the *m*/2 correlations is weighted based on the variation of the smoothed values at that distance such that the weights sum to one. Therefore, the resultant metric is restricted to [−1, 1] and has a similar interpretation as a standard Pearson correlation coefficient.

**Figure 1.**
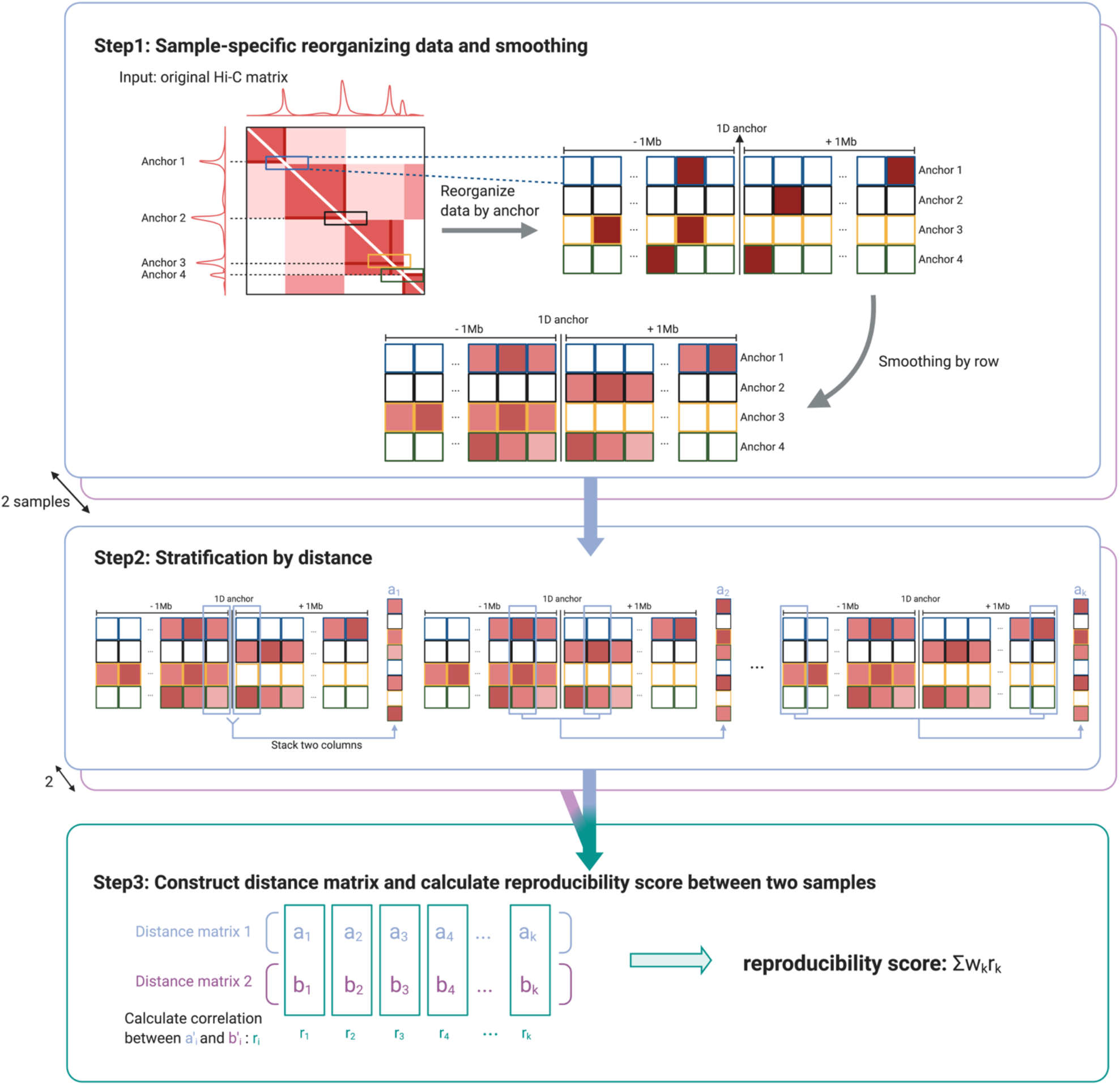
Cartoon illustration of HPRep. Step 1 involves first identifying anchors (i.e., 1D ChIP peak sites) and then extracting all interactions between these anchors and bins within a specified genomic distance from the anchors. This is followed by a one-dimensional smoothing procedure. Stratification by distance is performed in step 2 such that the elements of vector *a_k_* represent interactions that are equidistant from their respective anchors, *k* bins apart. In the final step, the Pearson correlation coefficients are calculated between *a_k_* from one sample and the analogous vector (*b_k_*) from the other sample for all *k*, and these Pearson correlation coefficients are combined in a weighted average to yield the final reproducibility metric.

Let *a*_*k*_ and *b*_*k*_ be two vectors of length 2*N* from samples *a* and *b* respectively, whose elements are normalized contact counts, where *N* represents the number of anchor bins in the union set of anchor bins from all samples in the study, and *k* indexes bins that are ±*k* units away. Let 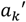 and 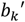 be the resulting vectors of length *N*_*k*_ ≤ 2*N* after removing any elements that are 0 in identical positions in both two vectors. The weight for stratum *k*, *w*_*k*_, is defined as

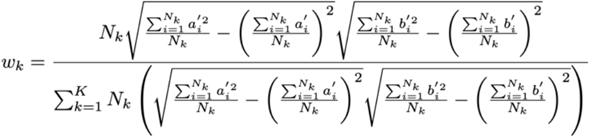

where *K* is the total number of strata, which is analogous to the weights used in HiCRep (Yang *et al.*, 2017). The numerator of *w*_*k*_ is the product of strata size and the standard deviations of 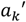 and 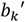, while the denominator is the sum of the numerators over all strata. Consequently, the weights are restricted to [0, 1] and sum to 1, where larger and more variable strata carry more weight than smaller and less variable strata.

## 3 Results

### 3.1 Mouse H3K4me3 PLAC-seq data

To evaluate the performance of HPRep, we first analyzed published H3K4me3 PLAC-seq datasets from mouse embryonic stem cells (mESCs) (Juric *et al.* 2019) and mouse brain tissues (Yamada *et al.* 2019), both consisting of two samples, by applying HPRep at 10Kb resolution. Samples from the same cell type or tissue were labeled as biological replicates while those cross cell type or tissue were labeled non-replicates, yielding two pairs of biological replicates and four pairs of non-replicates. Pseudo replicates were generated by pooling the two samples of the same cell type or tissue together, and then partitioning the pooled contact frequency in each bin pair randomly via Binomial (p = 0.5) sampling.

We would expect that pseudo replicates are most similar, followed by biological replicates, and that non-replicates are least similar. Indeed, this expected pattern is observed using HPRep (**Figure 2**), with results also exhibiting highly consistent patterns across chromosomes (**Supplementary Figure 1**). The higher metric for replicate mESC samples relative to mouse brain samples is due to the higher sampling depth of the former.

**Figure 2.**
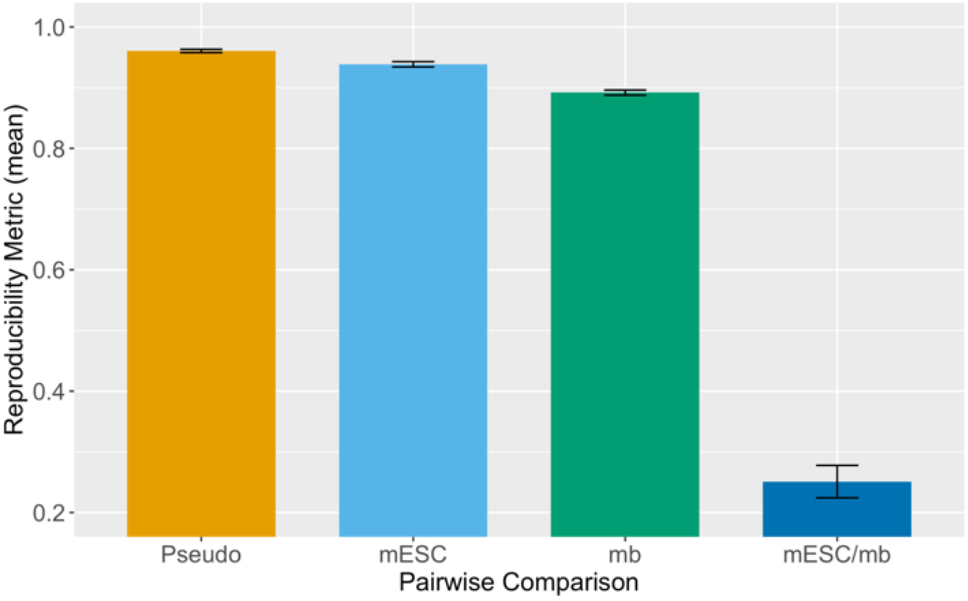
HPRep in mouse PLAC-seq data. Metrics obtained applying HPRep to PLAC-seq data from mESC and mouse brain (mb) tissues. Pseudo replicates generated from pooling two mESC samples followed by random sampling. Cross sample results represent mean of four pairings. Results are presented as the mean value over 19 autosomal chromosomes with error bar representing ± 1 standard deviation.

We next compared HPRep with alternative methods, specifically two Hi-C reproducibility methods: HiCRep (Yang *et al.*, 2017) and HiC-Spector (Yan *et al.*, 2017) as well as a naïve Pearson correlation (Supplementary Text S4). Since the Hi-C specific methods are designed using *n* × *n* symmetric contact matrices as the standard input, for these comparisons, in addition to restricting to bin pairs in the AND and XOR sets, we generated a “pseudo Hi-C” dataset from a HP dataset by also using all bin pairs (including the AND, XOR and NOT sets). The naïve Pearson correlation consisted simply of converting the entire upper triangular Hi-C contact matrices for each sample to single vectors and calculating the Pearson correlation coefficient between them. The methods were performed separately on all 19 autosomal chromosomes and the resulting metrics were reported as the arithmetic mean. The HiCRep and HiC-Spector methods were applied with the default parameters. The results are displayed in **Figure 3**.

**Figure 3.**
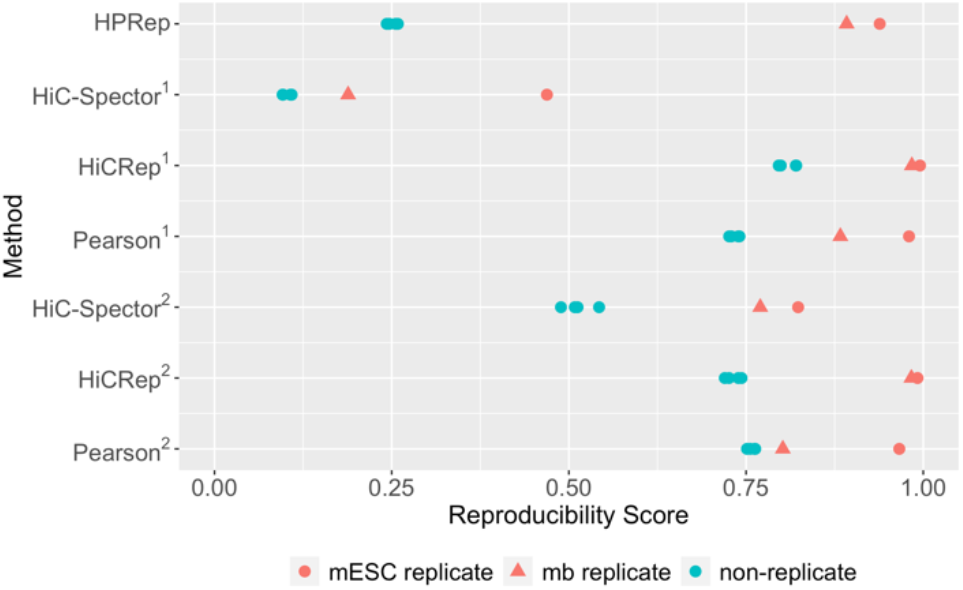
Comparison of methods, in mouse PLAC-seq datasets. HPRep compared to Hi-C specific methods HiC-Spector and HiCRep as well as Pearson correlation. 1: All methods using bin pairs in the AND and XOR sets. 2 Methods other than HPRep using all bin pairs in the AND, XOR and NOT sets. PLAC-seq dataset consisted of two mESC and two mouse brain replicates.

All methods except for naïve Pearson correlation yielded results consistent with what we expected, namely higher similarity for the biological replicates and lower similarity for the non-replicates. The similarity or reproducibility values for the biological replicates were similar among these three methods, which is expected for HPRep and HiCRep, since both methods are based on stratified Pearson correlation, but is noteworthy for HiC-Spector since it is based on a rather different method, and is restricted to [0, 1] as opposed to [-1, 1]. The difference among these methods, with the exclusion of HiC-Spector when including the NOT set, manifests largely in values for non-replicates, with HPRep yielding much smaller values relative to the others, although in each case the four non-replicate pair results were very consistent. Interestingly, the naïve Pearson correlation fails with the mouse brain sample, yielding a reproducibility score nearly identical to those of the non-replicates whereas the result from mESC replicates is consistent with the other three methods. This failure is obviated in HiCRep and HPRep, the other Pearson based methods. For example, for biological replicates, HPRep yields a mean reproducibility metric of 0.92 compared to a mean value of 0.25 for non-replicates. For the experiments using bin pairs in the AND, XOR and NOT sets, the mean reproducibility metrics comparing replicates and notn-replicates are 0.80 vs. 0.51, 0.99 vs. 0.73, and 0.88 vs. 0.76 for HiC-Spector, HiCRep, and Pearson correlation coefficients, respectively.

### 3.2 Human HiChIP data

In addition, we applied HPRep to measure the reproducibility of H3K27ac HiChIP data from GM12878 cells (two biological replicates) and K562 cells (three biological replicates) at 10Kb resolution (Mumbach *et al.*, 2017) resulting in 4 pairs of biological replicates (1 pair from GM12878, 3 pairs from K562) and 6 pairs of non-replicates (**Figure 4**). We anticipated *a priori* that differences between replicates and non-replicates would be more pronounced in this human dataset than the previous mouse H3K4me3 PLAC-seq dataset due to the greater dissimilarity in H3K27ac anchor bins between GM12878 cells and K562 cells. Specifically, the GM12878 and K562 cell lines contain 31,980 and 26,963 H3K27ac 10Kb anchor bins genome-wide (autosomal) respectively, with only 14,304 shared (Jaccard index 0.32). By contrast, mESC and mouse brain have 28,903 and 21,778 H3K4me3 10Kb anchor bins, with 17,722 overlapping, (Jaccard index 0.54) which is not expected since active promoters are largely shared across tissues and cell lines. For this human dataset, different methods were performed individually on all 22 autosomal chromosomes and the resulting metrics were averaged across chromosomes.

**Figure 4.**
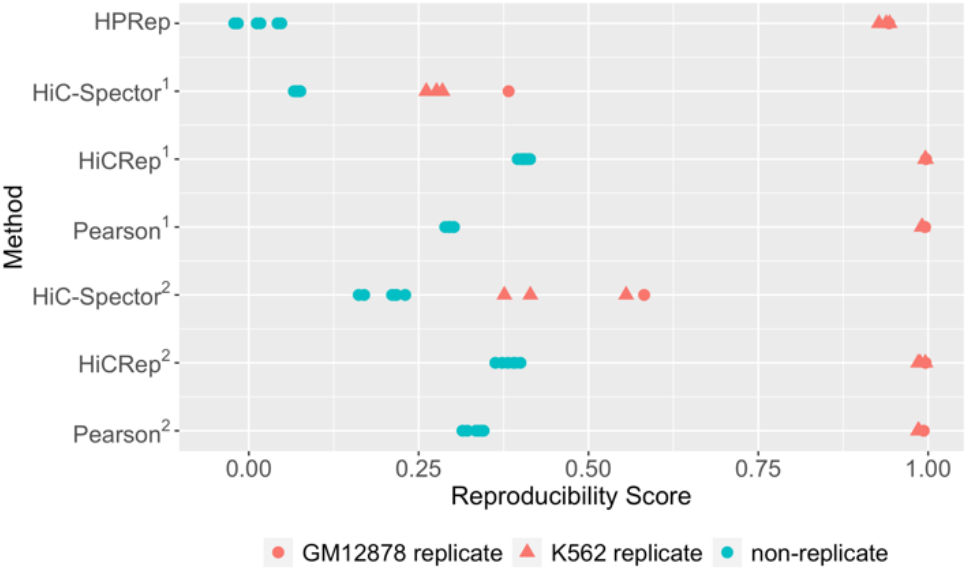
Comparison of methods in HiChIP datasets from human blood cell lines. HPRep compared to Hi-C specific methods HiC-Spector and HiCRep as well as Pearson correlation. ^1^ All methods using bin pairs in the AND and XOR sets. ^2^ Methods other than HPRep using all bin pairs in the AND, XOR and NOT sets. HiChIP dataset consisted of two GM12878 replicates and three K562 replicates.

The results from the human HiChIP data are consistent with those from mouse PLAC-seq data: the biological replicates yield high similarity (close to 1) while the non-replicates yield uniformly lower similarity. While all autosomal chromosomes were used in these analyses and results were largely consistent across them using HPRep, HiCRep, and Pearson correlation coefficients, results were quite inconsistent using HiC-Spector (**Supplementary Figure 2**). Specifically, HiC-Spector used 20 eigenvectors in the computation of a reproducibility metric, yet for several chromosomes convergence fails so fewer eigenvectors are used which yields erratic results (Supplementary Table 1). Again, HPRep results in the lowest metrics for the non-replicates which are all close to zero, highlighting the influence on anchor bin identity in this method.

### 3.3 Human PLAC-seq data

We next applied HPRep to a more complex H3K4me3 PLAC-seq dataset at 5Kb resolution, consisting of 11 samples from four brain cell types in human fetal brain obtained via fluorescence-activated cell sorting (Song *et al.* 2020): 3 samples from neurons (N), 3 samples from interneurons (IN), 2 samples from radial glial (RG), and 3 samples from intermediate progenitor cells (IPC). These samples have varying sequencing depths (detailed in Supplementary Table 2 of *Song et al.* 2020), with number of *cis* reads ranging from 47.5 million for RG2 (the second replicate of RG) to 390 million for RG1 (first replicate of RG). The anchor bins are defined as the union of 1D H3K4me3 peaks from all 4 cell types. In **Figure 5a**, reproducibility obtained by HiCRep shows no differentiation between inter- and intra-cells types. In contrast, HPRep shows a clear pattern of higher similarity for replicates from the same cell type compared to those from different cell types.

**Figure. 5.**
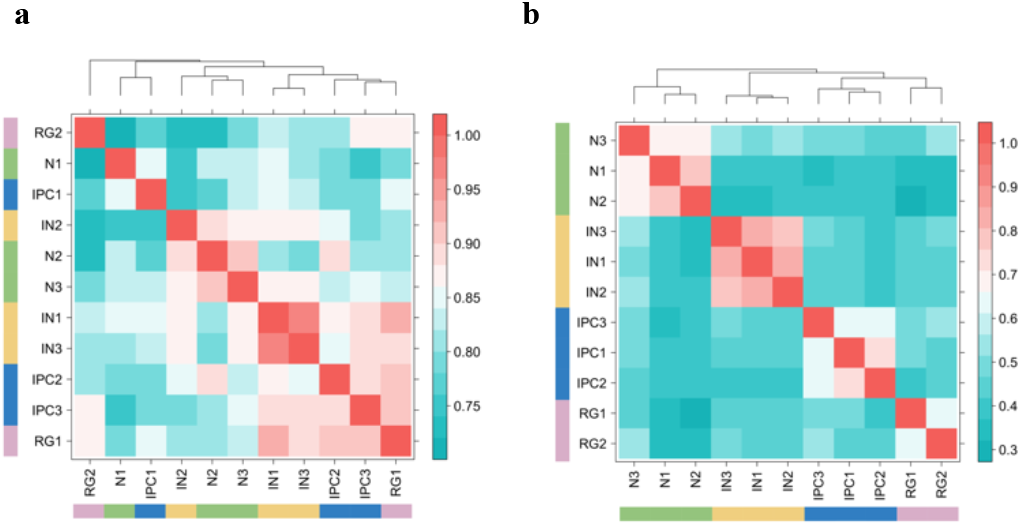
Comparison of HPRep and HiCRep in human brain PLAC-seq datasets. HPRep compared to HiCRep. HiChIP dataset consisted of 3 neuron, 3 interneuron, 2 radial glial, and 3 intermediate progenitor cell samples. Red color signifies results indicating stronger correlation.

Focusing on bin pairs in the AND and XOR sets highlights the effect of normalizing ChIP enrichment level. **Figure 6** is analogous to **5a** excluding bin pairs in the NOT set. The cell type clustering is more in line with the known truth, however, still has misspecifications according to the dendrogram: neuron, interneuron, and IPC cells are correctly grouped, but radial glial cells are misclassified into two groups.

**Figure. 6.**
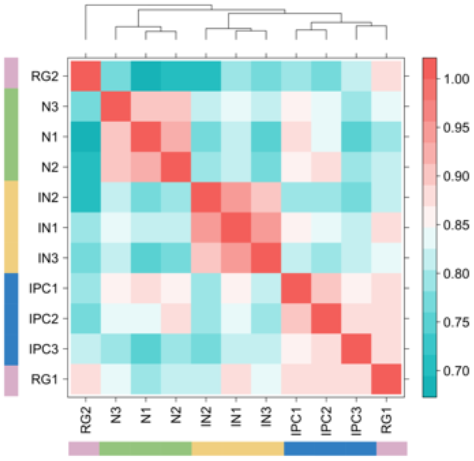
HiCRep excluding NOT pairs in human neural PLAC-seq datasets. HiCRep. HiChIP dataset consisted of three neuron, 3 interneuron, 2 radial glial, and 3 intermediate progenitor cell samples excluding interactions where neither bin overlapped with an anchor. Red color signifies results indicating stronger correlation.

Recent studies have shown that HiCRep is sensitive to sequencing depth (Yardimici *et al.*, 2019). To evaluate the robustness of HPrep with respective to different sequencing depths, we performed down-sampling to the original PLAC-seq data from 4 human brain cell types. This was performed by sampling from a multinomial distribution with *n* equal to the original count multiplied by a down-sampling factor and count probabilities set to match the distribution in the original data (Supplementary Text S5).

The first down-sampling was performed such that all samples matched the depth of the sample (RG2) which had the lowest sequencing depth. Note the identical color scales for **Figures 5b** and **Figure 7**, but the decrease in metric values for many pairwise comparisons for samples of the same cell type such as the interneuron cells. In order to quantify this reduced discernibility between samples, we utilized the silhouette procedure (Rousseeuw 1987), treating reproducibility score as a distance metric and reporting the average of the 11 silhouette values, one for each sample (Supplementary Text S6). We obtain 0.717 and 0.685 for the original experiment and down-sampled results respectively, where smaller numbers indicate worse clustering performance.

**Figure. 7.**
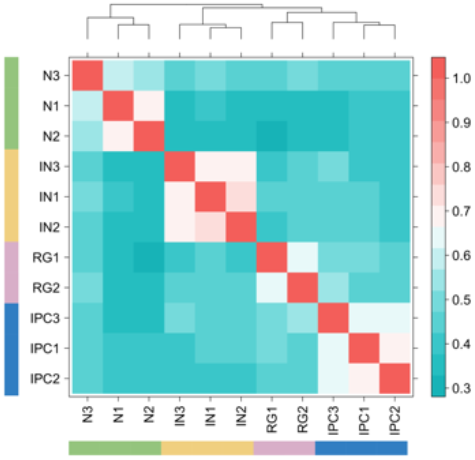
Performance of HPRep in down-sampled human neural PLAC-seq data. HPRep results obtained after down-sampling all eleven samples to read depth of the lowest sample.

Subsequent down-sampling was performed uniformly across all samples such that total counts were reduced to 80%, 60%, 40%, and 20% of their original values following the same sampling protocol as described above. As expected, in **Figure 8** we observe decreased discernibility among samples from different cell types, most strikingly with IPC and RG where the within sample HPRep reproducibility metric dropped to as low as 0.26 and 0.43, respectively. Applying the modified silhouette procedure described above to these four down-sampled datasets, we obtained a silhouette score of 0.700, 0.678, 0.634, and 0.518 for down-sampling to 80%, 60%, 40%, and 20% respectively.

**Figure. 8.**
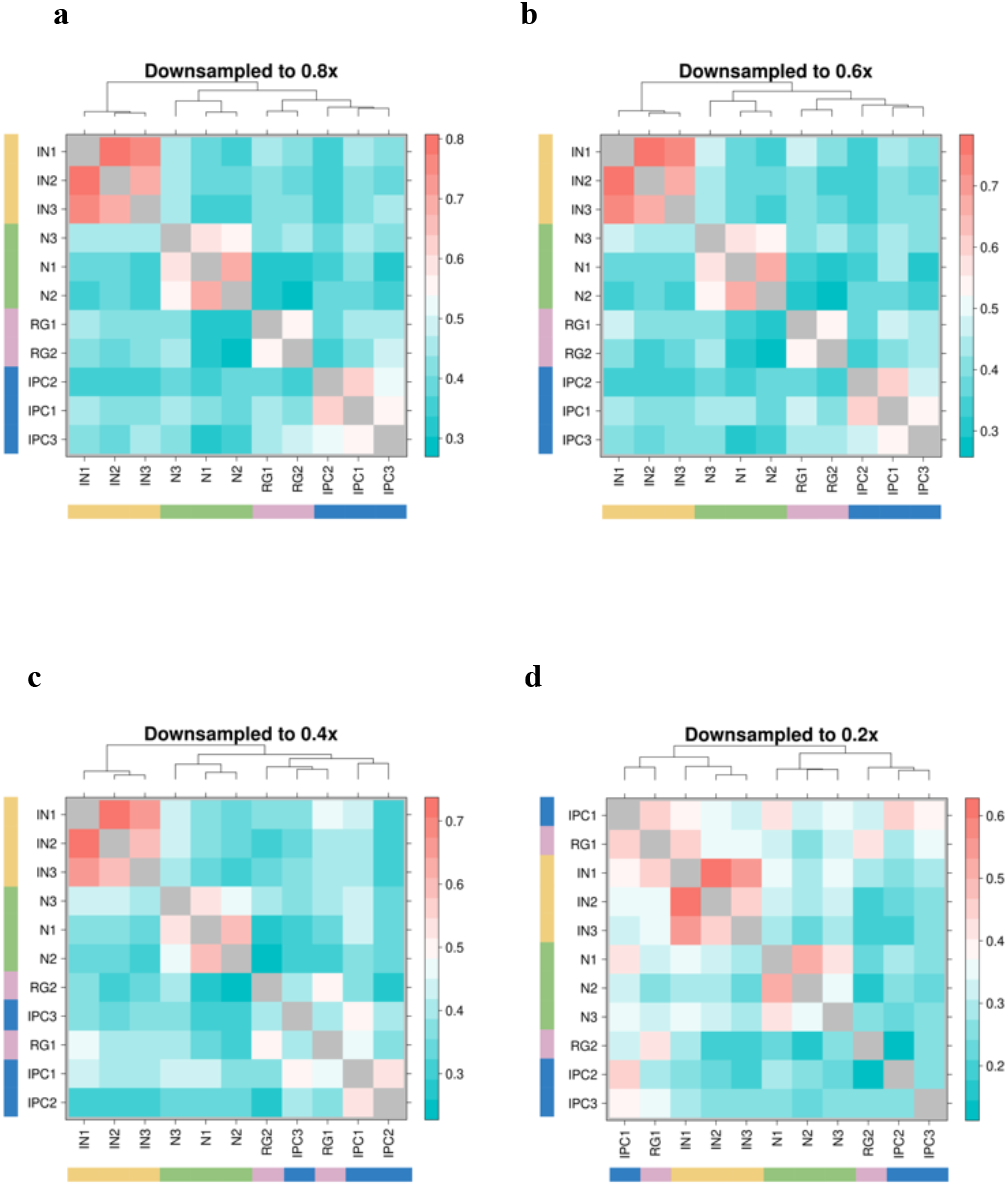
Down-sampling uniformly across all samples in human neural PLAC-seq datasets. HPRep results obtained after down-sampling each sample to by specified factor. a) 80% of original depth of each sample, b) 60% of original depth, c) 40% of original depth, d) 20% of original depth. Note that the diagonal is now gray to remove it from the scaling in order to better highlight difference.

## 4 Discussion

Quantification of data reproducibility is critical to ensure scientific rigor, yet methods tailored for HiChIP and PLAC-seq data are still lacking. Here, we propose HPRep, the first model-based approach to account for ChIP enrichment in measuring HP data reproducibility. Given the lack of HP specific tools, we compare HPRep to existing methods designed for Hi-C data, specifically HiCRep and HiC-Spector. Additionally, since our method, similar to HiCRep, relies on a weighted average of Pearson correlation coefficients, we also compare HPRep to the naïve Pearson correlation coefficient.

Our HPRep method, improving on existing Hi-C specific methods, is tailored to HP data for the measurement of reproducibility in two fundamental ways. First, HPRep is designed around the specific structure of HP data: while Hi-C data consists of contact frequencies among all bin pairs, HP data focuses on bin pairs where at least one bin overlaps with a ChIP-seq peak for a protein of interest. This is different from the standard *n* × *n* symmetric Hi-C contract matrix. We focus on the data matrix on anchor bins, regions that overlap with ChIP-seq peaks, and pairs between bins within a specified window of these anchors as illustrated in **Figure 1**.

Second, HPRep fits a positive Poisson regression model to normalize HP-specific ChIP enrichment and uses the residuals as the normalized contact frequencies. It also analyzes bin pairs in the AND and XOR sets separately, effectively accounting for ChIP enrichment for the two different types of bin pairs.

Our results from mouse H3K4me3 PLAC-seq data demonstrated very low variability in metrics between chromosomes (**Figure 1**), which is consistent with HiCRep (**Supplementary Figure 3**). In addition, we also compared HPRep with other existing methods using human H3K27ac HiChIP data from GM12878 and K562 cells, as well as H3K4me3 PLAC-seq data from 4 human brain cell types. Oure results demonstrated the superior performance of HPRep, in terms of accurate clustering of samples from the human brain cell types which was not achievable using HiCRep, although better clustering accuracy was observed when excluding bi pairs in the NOT set.

Future work involves exploring the potential of using this method to determine minimum per sample sequencing depth or maximum allowable (if any) differential depth across samples for accurate quantification of HP data reproducibility. We show that sample differentiation and expected clustering are robust to down-sampling, but rigorous analysis needs to be performed in order to demonstrate practical use, as more high-depth HP data become available from more tissues, cell lines or cell types. Additionally, we plan to examine the use of this general framework with capture Hi-C datasets, including those targeting at a relatively small number of loci identified from genome-wide association studies, and those genome-wide promoter capture Hi-C experiments. The use of pre-defined anchors by these methods encourages us to believe that the HPRep framework will be also applicable to these capture Hi-C methods, therefore these extensions are highly warranted but are beyond the scope of our current HPRep work.

In terms of computational efficiency, for the human PLAC-seq data consisting of 11 samples, tuning the smoothing parameter and determining all 55 pairwise reproducibility metrics for all 22 autosomal chromosomes took 1 hour and 5 minutes using a single core on a 2.50 GHz Intel processor with 4GB of RAM. One can choose to apply HPRep to one chromosome for almost same result. On the same data, HPRep takes 35 minutes to perform tuning and analysis on solely chromosome 1 using the same single core.

## 5 Conclusion

Here, we present HPRep, a computationally efficient algorithm based on positive Poisson regression (Juric *et al.*, 2019) and a stratified Pearson correlation (*Yang et al.*, 2017). Our comprehensive benchmark analyses of real HP datasets demonstrate that HPRep outperforms existing Hi-C reproducibility measurements.

## Acknowledgements

We would like to thank Dr. Di Wu for critical read of an earlier version of this manuscript. We also would like to thank the 4DN investigators for providing helpful comments to improve the utility of HPRep.

