## Supplementary Information for "HPRep: Quantifying reproducibility in HiChIP and PLAC-seq datasets"

**Text S1. Details for Step 1 and 2 of HPRep**

During the pre-processing step, intra-chromosomal reads are split into two groups: short-range reads (<= 1 Kb) and long-range reads (> 1Kb). The short-range reads are used as a measure of ChIP efficiency in the regression framework described later in the pipeline. Long-range reads are used to determine long-range interactions, which are extracted and classified as either AND, XOR, or NOT sets based on whether 2, 1, or 0 (respectively) read ends overlap with a ChIP-seq identified peak for the protein of interest. Additional details can be found in the MAPS paper: (Juric I. *et al*. (2019) MAPS: Model-based analysis of long-range chromatin interactions from PLAC-seq and HiChIP experiments. *PLOS Computational Biology*, 15(4): e1006982).

The regression and normalization step (step 2) follows a multi-step procedure:

1) We model the non-zero intra-chromosomal contacts as a zero-truncated Poisson model with mean $\mu_{ij}$. The covariates for effective fragment length (FL), GC content (GC), mappability (MS), and ChIP enrichment level (IP) are provided by the feather pre-processing step (as implemented in the MAPS pipeline), and represent $\log(x_i * x_j)$, where $x_i$ and $x_j$ are the corresponding covariate for bin *i* and *j* respectively. We fit regression models for the AND and XOR sets separately.

$$\log(\mu_{ij}) = \beta_0 + \beta_1 \cdot FL_{ij} + \beta_2 \cdot GC_{ij} + \beta_3 \cdot MS_{ij} + \beta_4 \cdot IP_{ij}$$

2) Fitted values are determined for each bin pair based on the resulting model for AND and XOR sets in each chromosome, resulting in 2 * n files where n is the number of autosomal chromosomes.
3) Normalized values are defined as $\log_2(1 + \text{observed} / \text{fitted})$ and all bin pairs are combined into one file. Additionally, the ChIP-seq peaks are binned to analysis resolution and supplied as a file containing a list of these anchor bins. Peaks that span a bin boundary are assigned to all bins they span.

**Text S2. Details for Step 3 of HPRep**

The final step involves data smoothing and sample comparison to calculate a final reproducibility metric between each pair of samples as a weighted Pearson correlation. The combined AND and XOR normalized data is stored in a matrix which is used as an input for the comparison algorithm. The basic data structure we consider is an $N$ x $m$ matrix, where $N$ represents the number of anchor bins in the union set of anchors from all samples and $m$ is 2 x binning distance/resolution, where binning distance is recommended to be set at 1Mb but can be user specified. Interactions further than 1Mb are typically sparse and highly variable. The $ij$ element of the matrix represents the normalized contact frequency between the anchor $i$ and the bin $j$ bin widths away, $j \in \{-m/2, ..., -1, 1, ... , m/2\}$. In the illustration below, $N = 4$, $m = 400$ at 5Kb resolution, 200 at 10Kb resolution.

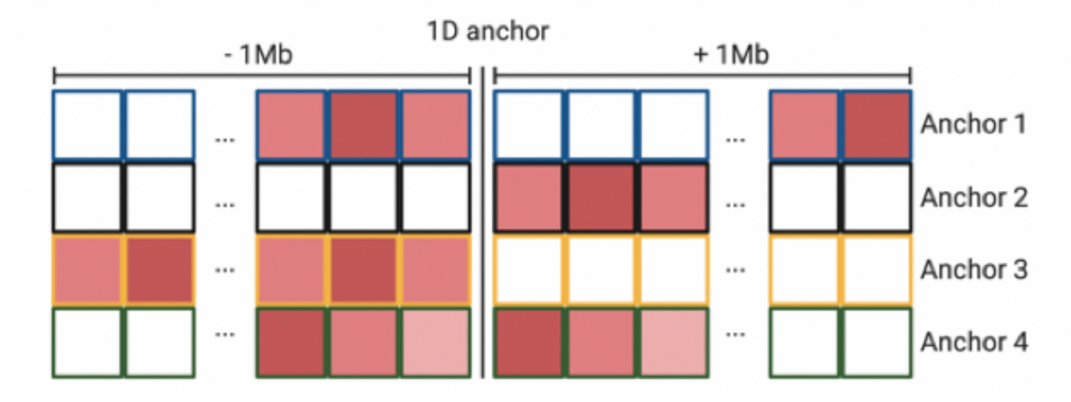


The normalized values undergo a 1-D smoothing procedure as follows: for a specified window size d, the *ij* element ($x_{ij}$) is transformed such that the smoothed value is

$$x_{ij}^{smoothed} = \left( \sum_{k\, =\, j-d}^{j+d} x_{ik} \right) \Big/ (2d+1)$$

Let $a_k$ and $b_k$ be two vectors of length 2*N* from samples *a* and *b* respectively, whose elements consist of the values from the smoothed data matrix from columns ± *k* units symmetrically from the center. All these values represent normalized and smoothed contacts that are ± *k* bins from their respective anchors. Let $a_k$' and $b_k$' be the resulting vectors of length $N_k <= 2N$ after removing any elements satisfying $a'_i = b'_i = 0$, where $a'_{ki}$ is the *i*th element of vector $a_k$'.

We define $r_k$ as

$$r_k = \frac{N_k \sum_{i=1}^{N_k} a'_i b'_i \; - \sum_{i=1}^{N_k} a'_i \sum_{i=1}^{N_k} b'_i}{\sqrt{N_k \sum_{i=1}^{N_k} a_i'^2 \; - \left(\sum_{i=1}^{N_k} a'_i\right)^2} \sqrt{N_k \sum_{i=1}^{N_k} b_i'^2 \; - \left(\sum_{i=1}^{N_k} b'_i\right)^2}}$$

namely the empirical correlation between $a'_k$ and $b'_{k.}$ The define the weights for each of the k strata as

$$w_k = \frac{N_k \sqrt{\frac{\sum_{i=1}^{N_k} a_i'^2}{N_k} - \left(\frac{\sum_{i=1}^{N_k} a_i'^2}{N_k}\right)^2} \sqrt{\frac{\sum_{i=1}^{N_k} b_i'^2}{N_k} - \left(\frac{\sum_{i=1}^{N_k} b_i'^2}{N_k}\right)^2}}{\sum_{k=1}^{K} N_k \left(\sqrt{\frac{\sum_{i=1}^{N_k} a_i'^2}{N_k} - \left(\frac{\sum_{i=1}^{N_k} a_i'^2}{N_k}\right)^2} \sqrt{\frac{\sum_{i=1}^{N_k} b_i'^2}{N_k} - \left(\frac{\sum_{i=1}^{N_k} b_i'^2}{N_k}\right)^2}\right)}$$

The reproducibility score between two matrices is then the weighted average of the stratified correlations $r_k$

$$\text{reproducibility score} = \sum_{k=1}^{K} r_k w_k$$

**Text S3. Smoothing parameter optimization**

The smoothing parameter d is tuned using the method similar to the HiCRep protocol with modification to the sampling scheme and search termination criterion. The following algorithm is used:

Two samples to be analyzed are selected, preferably ones that are dissimilar such as non-biological replicates. Twenty-five percent of the non-zero contacts from one are randomly sampled and used to populate a contact matrix as previously diagrammed, with the remaining entries set to zero. The analogous positions in the other sample are used to populate a corresponding matrix. The reproducibility score is calculated for these matrices and the sampling procedure is repeated a total of ten times with no smoothing performed. The average of these ten values is recorded.

The smoothing parameter is then iterated, repeating the above procedure until the average metric using smoothing parameter d+1 compared to d exhibits less than a one percent increase. The value of d is recorded and used as the smoothing parameter for all analyses with the particular dataset.

**Text S4. Procedures for comparative methods**

HiCRep:

All results obtained using HiCRep were conducted using R (3.6.0) and using version 1.12.0 of the HiCRep package obtained from https://github.com/MonkeyLB/hicrep. Default parameters were used for all experiments. Note that the documentation recommends a smoothing parameter of 20 for 10kb resolution but does not specify a recommended parameter for 5kb resolution. We used 20 for 5kb as well since marginal difference was reported when tuning beyond 20.

To ensure proper data formatting for use with HiCRep, the built-in function “bed2mat” was utilized, which converts a 3-column contact matrix to a square contact matrix with all elements not supplied set to 0. Experiments that included solely AND and XOR sets of contacts were prepared by extracting bin pairs and observed (integer) contacts from the corresponding AND / XOR files and those that also included NOT sets were generated similarly.

HiC-Spector:

The Python version of HiC-Spector was used rather than the Julia version since the former readily accepts Hi-C data in genomic coordinates rather than .hic format. The program used was “run_reproducibility_v2.py” found at https://github.com/gersteinlab/HiC-spector. Experiments that included solely AND and XOR sets of contacts were prepared by extracting bin pairs and observed (integer) contacts from the corresponding AND / XOR files. Note, the bin positions had to be converted to indices starting at 1, so the global minimum bin position was determined, and all bins positions scaled by (genomic position – minimum position) / resolution. Experiments also including NOT sets were generated similarly.

Pearson correlation:

The upper triangular component of a standard symmetric n x n contact matrix was flattened to a vector for each sample. The Pearson correlation between two samples was computed as the correlation between these vectors.

**Text S5. Down-sampling procedure**

The generalized down-sampling procedure was performed on the AND and XOR contact files for each chromosome separately. Let n be the total number of counts for all bin pairs in the specific file and let d be the down-sampling coefficient. That is, to down-sample to 0.8x depth, d = 0.8. The vector v of counts for all bin pairs is down-sampled to depth d utilizing the R function “rbinom” where the size parameter is set to floor(n * d) and the probability vector is the element-wise division of v by n. These down-sampled AND and XOR files then intersect the pipeline as usual with the removal of bins that now have counts of 0.

**Text S6. Determination of Silhouette Values**

Silhouette values were calculated via the method in Rousseeuw (1987). Let *d*(*i, j*) be the similarity between sample *i* and *j*, which is this analysis is the scaled reproducibility metric between the two samples. The silhouette method requires that the similarity (or distance) quantities be comparable on a ratio scale, that is, if the distance between two points is doubled that implies the points are twice as far apart. Pearson correlation does not have such a property, so for each experiment the values were standardized to [0, 1] by subtracting the lowest value and dividing by (max – min) value.

Let sample *i* be a member of cluster *A*. Furthermore, let *a(i)* be the average similarity of *i* to all other samples in the same cluster. Let *d(i, C)* be the average similarity of sample *i* to all other samples in cluster *C* and let *b(i)* be the maximum value of *d(i, C)* over all clusters C distinct from cluster *A.* Then the silhouette value is defined as

$$s(i) = \frac{a(i) - b(i)}{\max\{a(i), b(i)\}}$$

We report the average *s(i)* over all 11 samples. The closer this value is to 1 the better the clustering performance.

**Text S7. Data details**

For the human brain PLAC-seq data, fastp (https://github.com/OpenGene/fastp) was used to trim the fastq files to 100bp. No additional modifications to the described pipeline were performed on any of the datasets used in this paper. Default software options described in https://github.com/yunliUNC/HPRep were used for alignment and merging for all samples analyzed. Resolutions used for each dataset were:

| | |
|---|---|
| Mouse embryonic stem cell and mouse brain tissue H3K4me3 PLAC-seq: | 10Kb |
| Human brain H3K4me3 PLAC-seq: | 5Kb |
| GM12878 and K562 H3K27ac HiChIP: | 10Kb |

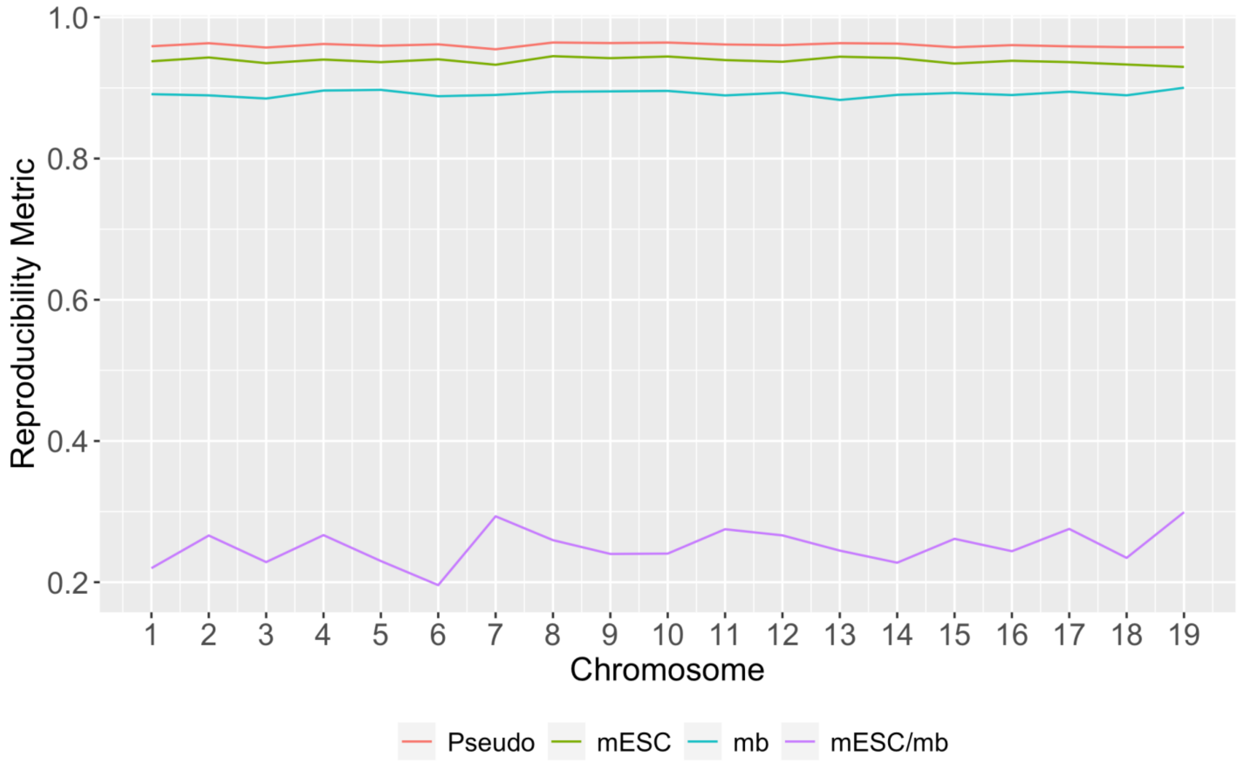


**Supplementary Fig. 1 Reproducibility metric by chromosome**.
Metrics obtained applying HPRep to two mouse embryonic stem cell (mESC) and mouse brain (mb) tissues H3K4me3 PLAC-seq samples. Pseudo replicates were generated from pooling mESC samples followed by random sampling via a Binomial (p=0.5) distribution. Cross sample results represent the mean of four cross-tissue pairings.

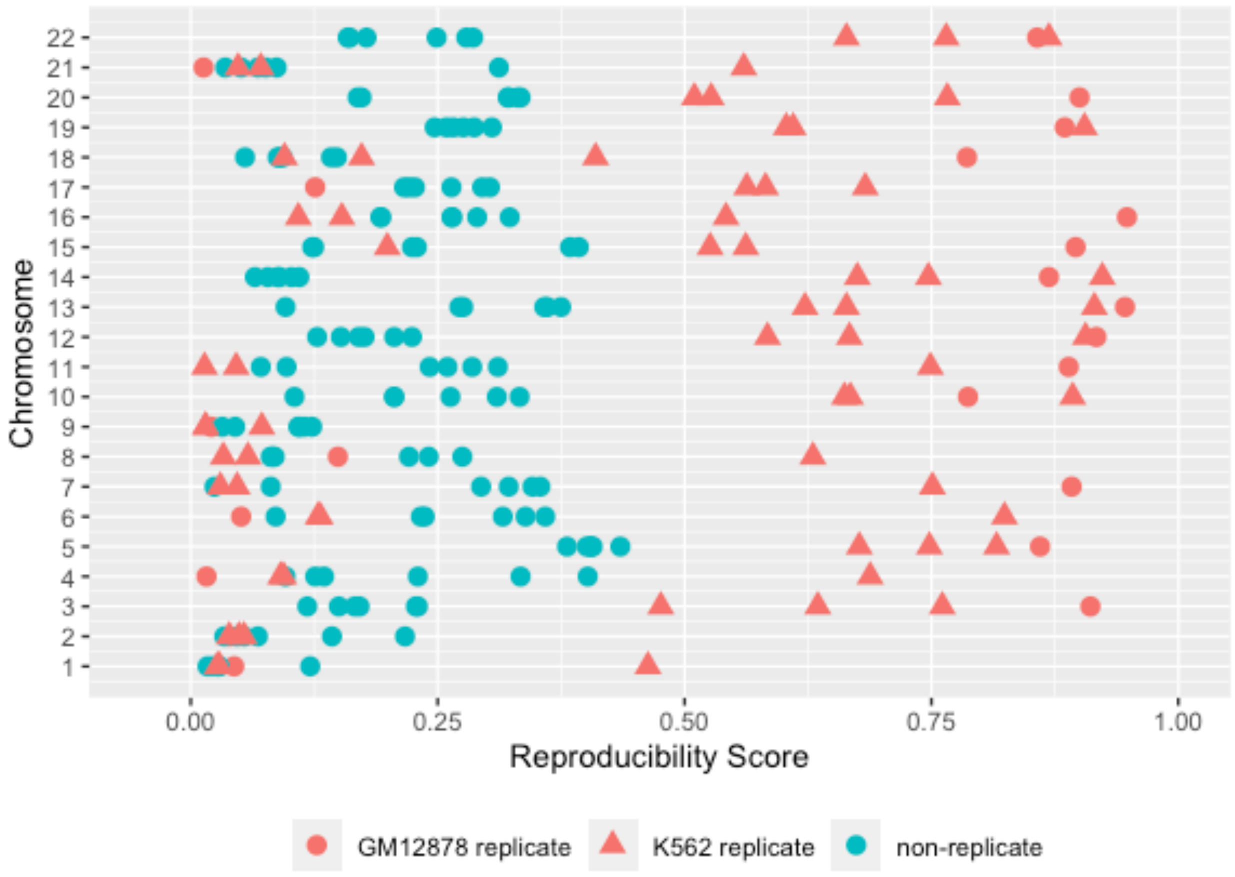


**Supplementary Fig. 2 HiC-Spector results by chromosome**. The plotted results clearly demonstrate the chromosome to chromosome variability we do not see with HiCRep or HPRep with this data. For example, the chromosome 22 results are as expected whereas the chromosome 21 results fail to distinguish between 5 of the 6 non-replicates and 3 of the 4 replicates.

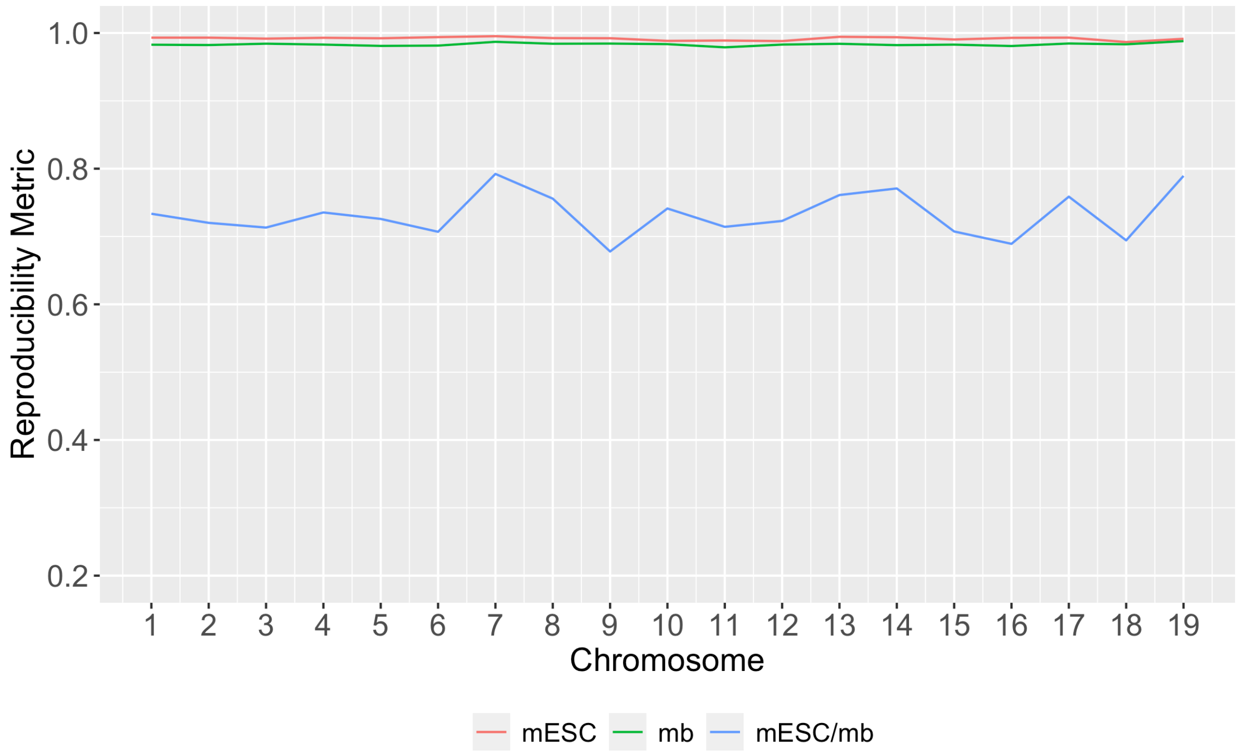


**Supplementary Fig. 3 HiCRep reproducibility metric by chromosome**.
Metrics obtained applying HiCRep to two mouse embryonic stem cell (mESC) and mouse brain (mb) tissues H3K4me3 PLAC-seq samples. Cross sample results represent the mean of four cross-tissue pairings.

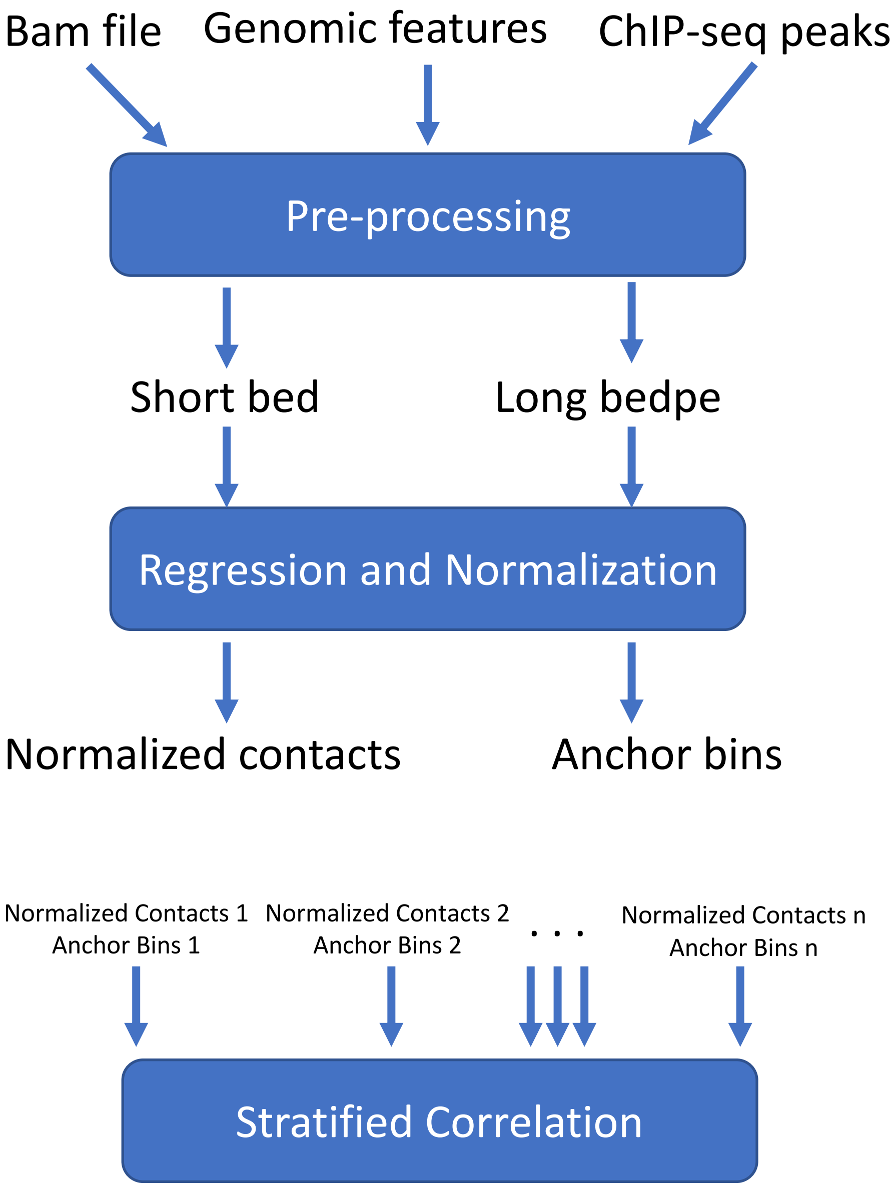


**Supplementary Fig. 3. HPRep workflow**

**Supplementary Table 1. Number of effective eigenvectors in HiC-Spector analysis**

| | Chromosome | | | | | | | | | | | | | | | | | | | | | |
|---|---|---|---|---|---|---|---|---|---|---|---|---|---|---|---|---|---|---|---|---|---|---|
| | 1 | 2 | 3 | 4 | 5 | 6 | 7 | 8 | 9 | 10 | 11 | 12 | 13 | 14 | 15 | 16 | 17 | 18 | 19 | 20 | 21 | 22 |
| GM12878 replicate | 18 | 19 | 20 | 18 | 20 | 19 | 20 | 20 | 18 | 19 | 20 | 20 | 20 | 19 | 20 | 20 | 19 | 19 | 20 | 20 | 16 | 19 |
| Non-replicate 1 | 19 | 19 | 20 | 19 | 20 | 19 | 20 | 18 | 19 | 19 | 20 | 20 | 20 | 19 | 20 | 19 | 19 | 20 | 20 | 20 | 16 | 20 |
| Non-replicate 2 | 18 | 19 | 20 | 19 | 20 | 19 | 18 | 20 | 19 | 20 | 18 | 20 | 20 | 19 | 19 | 19 | 19 | 19 | 20 | 20 | 16 | 20 |
| Non-replicate 3 | 19 | 19 | 20 | 18 | 20 | 19 | 19 | 20 | 19 | 20 | 20 | 20 | 20 | 19 | 20 | 20 | 19 | 19 | 20 | 20 | 16 | 20 |
| Non-replicate 4 | 17 | 19 | 20 | 18 | 20 | 19 | 20 | 18 | 19 | 19 | 20 | 20 | 19 | 20 | 20 | 19 | 19 | 18 | 20 | 20 | 18 | 20 |
| Non-replicate 5 | 17 | 20 | 20 | 18 | 20 | 18 | 18 | 20 | 19 | 19 | 18 | 20 | 20 | 20 | 19 | 19 | 19 | 20 | 20 | 20 | 18 | 20 |
| Non-replicate 6 | 17 | 18 | 20 | 18 | 20 | 19 | 19 | 20 | 19 | 19 | 20 | 20 | 20 | 19 | 20 | 20 | 19 | 20 | 20 | 20 | 18 | 20 |
| K562 replicate 1 | 16 | 19 | 20 | 19 | 20 | 19 | 19 | 20 | 19 | 19 | 18 | 20 | 19 | 20 | 18 | 19 | 20 | 18 | 20 | 20 | 19 | 19 |
| K562 replicate 2 | 19 | 19 | 20 | 19 | 20 | 19 | 19 | 20 | 20 | 19 | 20 | 20 | 19 | 19 | 20 | 20 | 20 | 18 | 20 | 20 | 19 | 19 |
| K562 replicate 3 | 18 | 18 | 20 | 19 | 20 | 19 | 19 | 20 | 19 | 20 | 19 | 20 | 20 | 19 | 18 | 19 | 20 | 18 | 20 | 20 | 18 | 20 |

Table 1 displays the number of effective eigenvectors used in the analysis of the cohesion HiChIP dataset using NOT data from Figure 3. In conjunction with Supplemental Figure 1 it illustrates how utilization of fewer than 20 eigenvectors affect the outcome with respect to expected values. For example, the analyses of chromosomes 3 and 12 utilize 20 eigenvectors for all 10 pairs and the biological replicates all have higher metrics than the non-replicates. Compare this with the results for chromosomes 1 and 2, whose analyses utilize almost exclusively fewer than 20 eigenvectors and the biological and non-replicates are not distinguished.

**Supplementary Table 2. Data sources**

| Data Description | Reference (PMID) | GEO accession number or other sources |
|---|---|---|
| mESC H3K4me3 PLAC-seq | 30986246 | GSE119663 |
| Mouse brain H3K4me3 PLAC-seq | 31068695 | GSE127995 |
| GM12878 H3K27ac HiChIP | 28945252 | GSE101498 |
| K562 H3K27ac HiChIP | 28945252 | GSE101498 |
| Human brain H3K4me3 PLAC-seq | 33057195 | Neuroscience Multi-Omic Archive (NeMO Archive) under controlled access |
